# Local transcriptional covariation produces accurate estimates of cell phenotype

**DOI:** 10.1101/2023.11.09.566448

**Authors:** Sinan Ozbay, Aditya Parekh, Rohit Singh

## Abstract

The utility of single-cell RNA sequencing (scRNA-seq) is premised on the notion that transcriptional state can faithfully reflect cell phenotype. However, scRNA-seq measurements are noisy and sparse, with individual transcript counts showing limited correlation with cell phenotype markers such as protein expression. To better characterize cell states from scRNA-seq data, researchers analyze gene programs–—sets of covarying genes–—rather than individual transcripts. We hypothesized that more accurate estimation of gene covariation, especially at a local (i.e., cell-state) rather than global (i.e., experimental) scale, could better capture cell phenotypes. However, the field lacks appropriate mathematical frameworks for analyzing gene covariation: coexpression is quantified as a symmetric positive-definite matrix, where even basic operations like arithmetic differences lack biological interpretability. Here we introduce Sceodesic, which exploits the Riemannian manifold structure of gene coexpression matrices to quantify cell state-specific coexpression patterns using the log-Euclidean metric from differential geometry. Unlike principal components analysis and non-negative matrix factorization, which infer only global covariation, Sceodesic efficiently discovers local covariation patterns and organizes them into interpretable, linear gene programs. Sceodesic outperforms existing approaches in predicting protein expression levels, distinguishing transcriptional responses to gene perturbations, and identifying biologically meaningful programs in fetal development. By respecting the mathematical structure of gene coexpression, Sceodesic bridges the gap between biological variability and statistical analysis of scRNA-seq data, enabling more accurate characterization of cell phenotypes.

**Software availability:** https://singhlab.net/Sceodesic

---

Single-cell RNA sequencing (scRNA-seq) has revolutionized our understanding of cellular heterogeneity. Fundamental to its applicability is the expectation that a cell’s transcriptome serves as an accurate proxy for its phenotype. As a close corollary, biological variation, whether temporal, perturbational, cross-modal, or otherwise, is expected to be reflected in meaningful transcriptional signatures. While individual marker genes sometimes indicate cell types or high-level phenotypes, cell types and states are more often characterized by coordinated expression of multiple genes, termed gene *programs*. Given scRNA-seq data’s inherent noise, robust identification of these programs is a critical challenge in generating informative transcriptional signatures (Fig. 1A).

**Figure 1:**
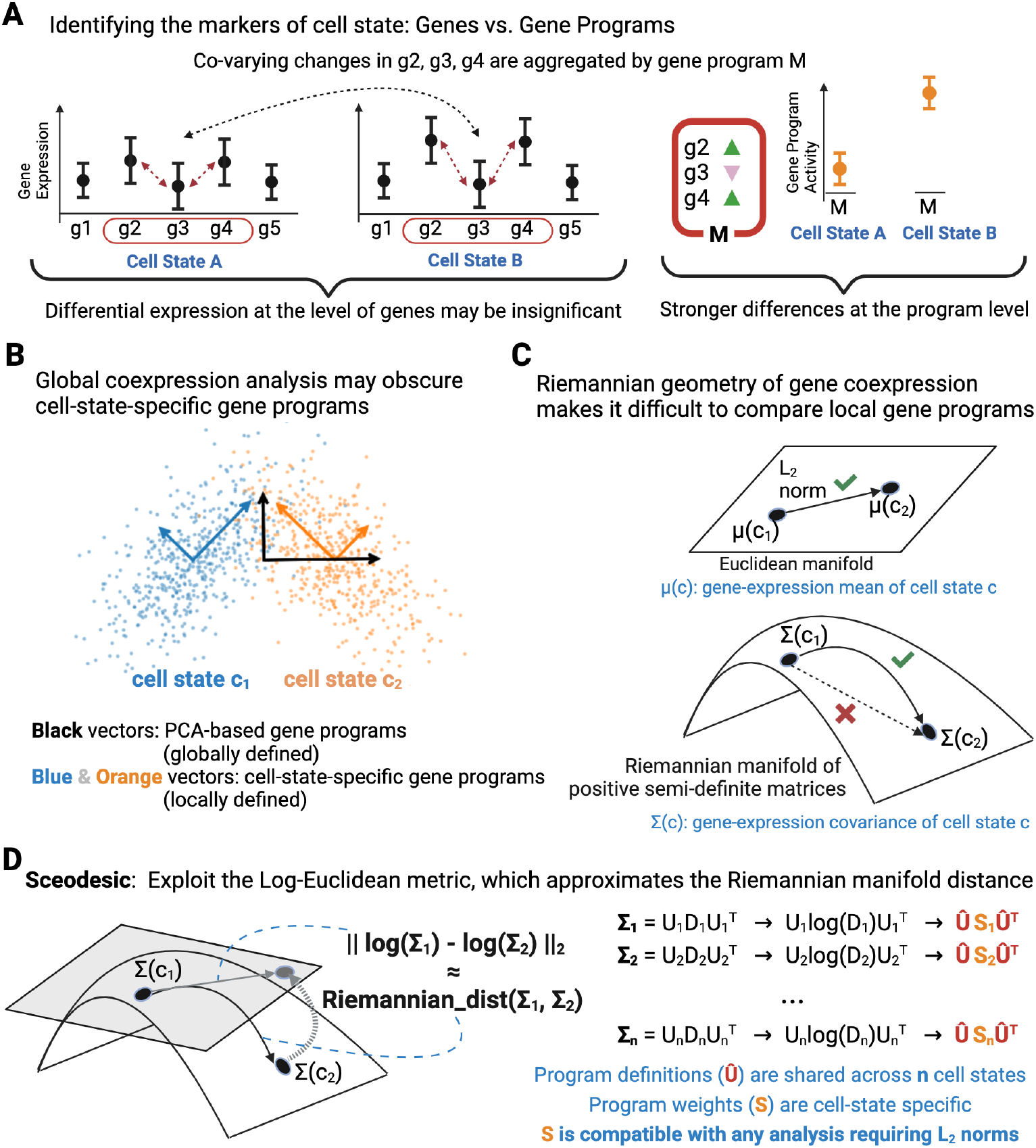
Leveraging Riemannian geometry for discovering scRNA-seq gene programs. **A**) Transcriptional drivers of cell differentiation and fate commitment may be more visible at the scale of gene programs (sets of co-varying genes) rather than individual genes since the former aggregate information over many genes. **B**) Global techniques like PCA are unable to access state-specific variations. With large scRNA-seq datasets now available, subsets of the data (cell cohorts) can be analyzed to uncover cell state-specific gene programs. **C**) Gene covariance obeys Riemannian, not Euclidean, geometry making it a challenge to compare and score gene programs with naive Euclidean distances. **D**) Sceodesic addresses this by approximating the Riemannian manifold distance with the Log-Euclidean metric, defined on the tangent bundle of the manifold. Using sparse eigenanalysis, we aggregate locally-defined programs and construct programs whose definitions are consistent across cell states and state-specific weights are amenable to Euclidean distances.

As linear, unsupervised approaches for representing combinations of genes, principal components analysis (PCA) and non-negative matrix factorization (NMF) have long been popular in scRNA-seq analysis. Their linearity ensures clear interpretability of each gene’s role, facilitating follow-up experiments like knockouts. Such approaches also uncover innate transcriptomic structure independent of specific biological covariates, thus reducing the risk of over-fitting or discovering spurious relationships. Moreover, these unsupervised methods work well with limited or noisy data, which has been an important concern, particularly in earlier scRNA-seq studies.

However, with technological advances, single-cell datasets have grown dramatically and now regularly profile millions, rather than hundreds or thousands, of cells. Simultaneously, independent measurements from different data modalities are increasingly available. These developments present both the opportunity and necessity to revisit the estimation of gene programs. The *opportunity* lies in using multimodal data to objectively assess the biological informativeness of estimated programs. For instance, Ji et al. [1] have benchmarked different metrics on scRNA-seq data using independently measured biological covariates. While these benchmarks enable evaluating the correlation between a distance metric applied to single cell data and biological covariates, correctly tracking transcriptomic differences between two cell states is often insufficient for a metric to deliver useful biological insights. Rather, it is typically also necessary to identify the *sets* of genes playing a dominant role in explaining such differences, which is precisely what program estimation methods do. The opportunity is to generate novel, informative transcriptional covariates that eventually guide experiments. The *necessity* to revisit gene program estimation arises because PCA and NMF become less suitable for larger, more heterogeneous datasets spanning over hundreds of cell types and dozens of conditions. Since these methods estimate covariance structures across the *entire* scRNA-seq dataset, they risk conflating broad technical artifacts with true biological variation, resulting in misleading or biologically irrelevant interpretations. For instance, consider the Simpson’s paradox-like scenario in Figure 1B. Although cell states *c*_1_ and *c*_2_ in the figure have the same local, state-specific principal components (blue and orange arrows), the standard or global principal components analysis (black arrows) completely misses those same cell state-specific directions of variation and thus resolves the data into gene programs that are not relevant for either cell state.

Fortunately, estimating *local* gene-gene coexpression structures can address the pitfalls of existing methods in dealing with complex modern scRNA-seq datasets. However, a mathematically rigorous, biologically faithful, and computationally efficient abstraction for local gene coexpression has been elusive. While arithmetic differences in expression levels are meaningful for individual genes, this approach fails for covariance matrices, which have a special mathematical structure. The values in these matrices are constrained by each other: if the gene pairs (A, B) and (B, C) each demonstrate high positive correlations, then the correlation between (A, C) cannot be very negative. Mathematically, these constraints imply that a covariance matrix must be positive semi-definite (PSD) and thus does not adhere to Euclidean geometry. To quantify coordinated changes in gene expression, we need to compare covariance matrices across these changes. But naively subtracting one covariance matrix from another ignores the curvature in the manifold of PSD matrices, risking inconsistent representations of transcriptional states (Fig. 1C). This is analogous to the observation that the shortest path between two cities on the Earth’s surface (e.g., the flight path between New York and Paris) does *not* correspond to a straight line between the two cities on the flattened Mercator map.

Our central observation is that the availability of large scRNA-seq datasets now enables the tools of differential geometry to be employed for principled, quantitative analyses of locally-defined covariance patterns. To that end, our core conceptual advance is to introduce the logarithmic map on the Riemannian manifold of positive semi-definite matrices, developing a mathematically consistent and biologically relevant metric for gene coexpression (Fig. 1D). This log-Euclidean metric is the first to respect the curvature of the covariance matrix space while preserving gene-covariance semantics. In single-cell analysis, this advance provides the first mathematically rigorous, linear, interpretable, and computationally practical means of comparing complex differences between cellular states based only on each cell state’s covariance matrix.

Building on this mathematical foundation, we present Sceodesic, an algorithm that combines differential geometry and spectral analysis to extract interpretable and cell state-specific gene expression programs from scRNA-seq data (Fig. 1D). Sceodesic takes a bottom-up approach: First, Sceodesic groups cells with similar transcriptional profiles into cohorts roughly corresponding to cell state, then estimates local covariance patterns and a set of gene programs for each of these cohorts. Second, from these local dictionaries, it learns a single, study-wide gene program dictionary and then reconstructs local covariance using the study-wide dictionary with the log-Euclidean metric as the objective function. The result is a single set of informative gene programs and a recapitulation of gene expression data as program expression that is both low-dimensional and captures information difficult to discern via individual genes, top-down estimation methods, or supervised approaches.

We validate Sceodesic against ground-truth biological data, comparing its ability to recapitulate protein expression (CITE-seq), perturbation state (Perturb-seq), and developmental trajectory better than PCA or NMF. Sceodesic excels in generating biologically meaningful and accurate gene program signatures for various sources of biological variation (Figs. 2, 3, 4, 5). It appends to each cell’s representation a feature vector encapsulating cell-state-specific gene programs, relying solely on covariance rather than mean expression levels. By introducing deep mathematical principles into a practical framework, Sceodesic is poised to drive fundamental and translational insights in single-cell transcriptomics and beyond.

**Figure 2:**
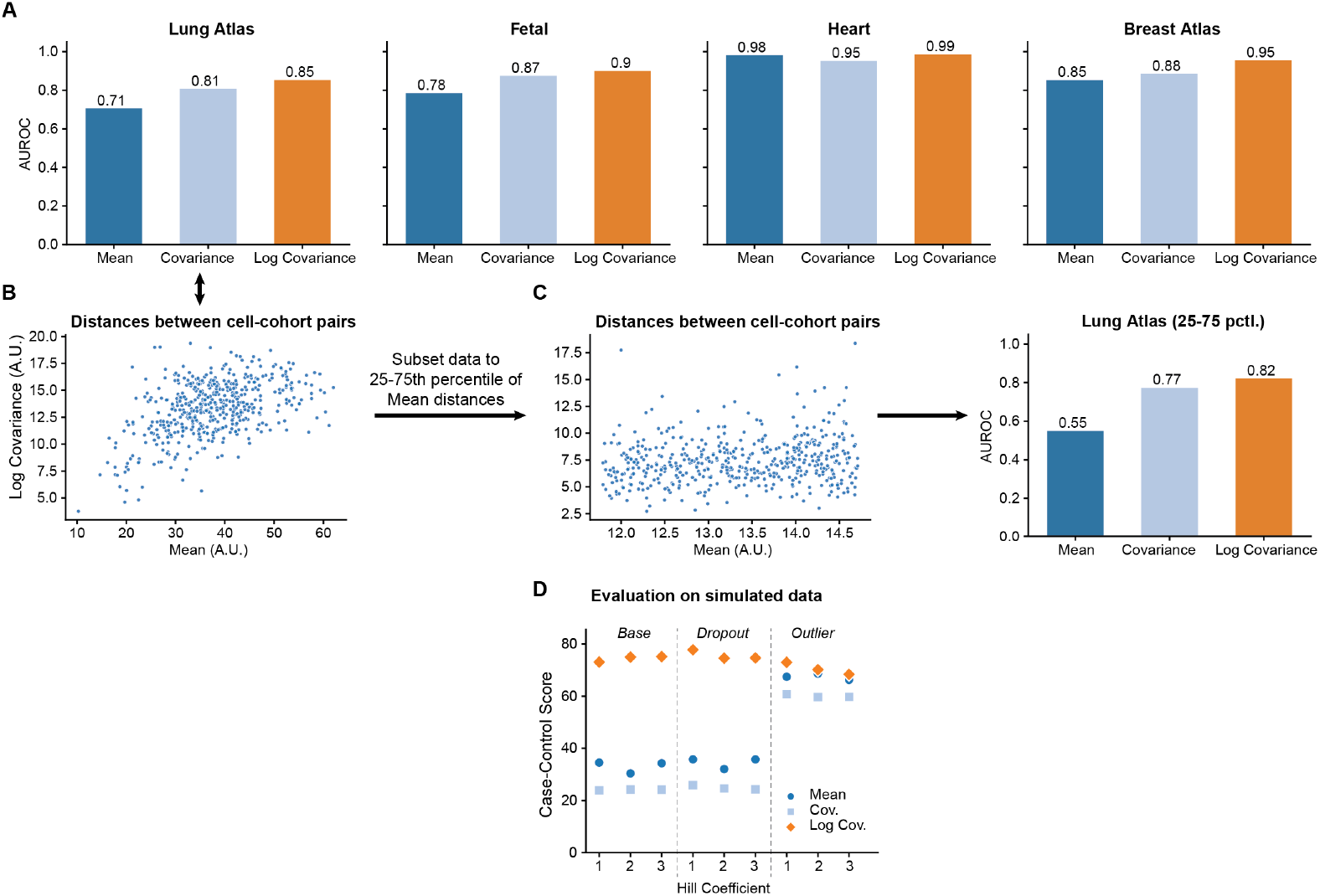
Comparison of distance metrics between cell cohorts: **A)** Comparison of Euclidean distances between cohort means (*dµ*), Euclidean distance between covariance matrices (*d*_Σ_), and log-Euclidean distances between covariance matrices (*d*_log(Σ)_) as 1-D classifiers of cohort-cell type similarity as measured by AUROC. **B)** Points signify cohort pairs with position (*d*_*µ*_, *d*_log(Σ)_), showing a stronger relationship between the mean and log-covariance at small distances. **C)** Identical analysis restricted to middle quartiles on the lung dataset show a weaker relationship of *d*_*µ*_ and *d*_log(Σ)_ on medium distances, with classifier outperformance persisting. (A.U. = Arbitrary Units) **D)** Comparison of the same distance metrics on simulated scRNA-seq data with added noise. case–control score is the ratio of the inter-cohort distance to the intra-cohort distance.

**Figure 3:**
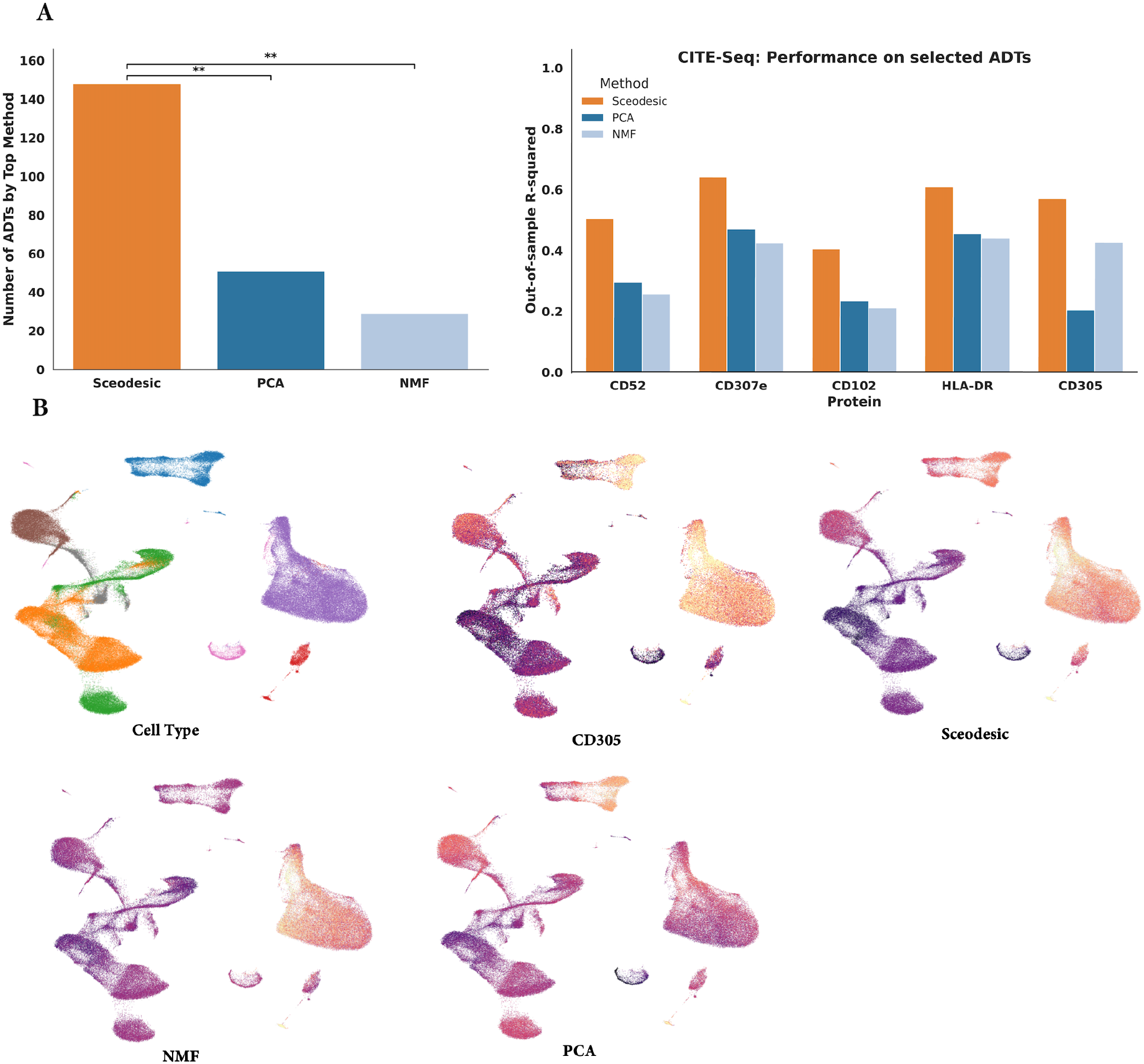
Comparing program estimation methods on CITE-Seq data. **A)** Number of ADT surface proteins whose expression levels are best predicted from scRNA-seq data resolved into Sceodesic, PCA, and NMF gene programs, as measured by out-of-sample R-squared. For each program estimation method and each surface protein, a Lasso regression was trained to assess program predictiveness. **B)** Performance of program estimation methods as measured by out-of-sample R-squared on selected proteins.

**Figure 4:**
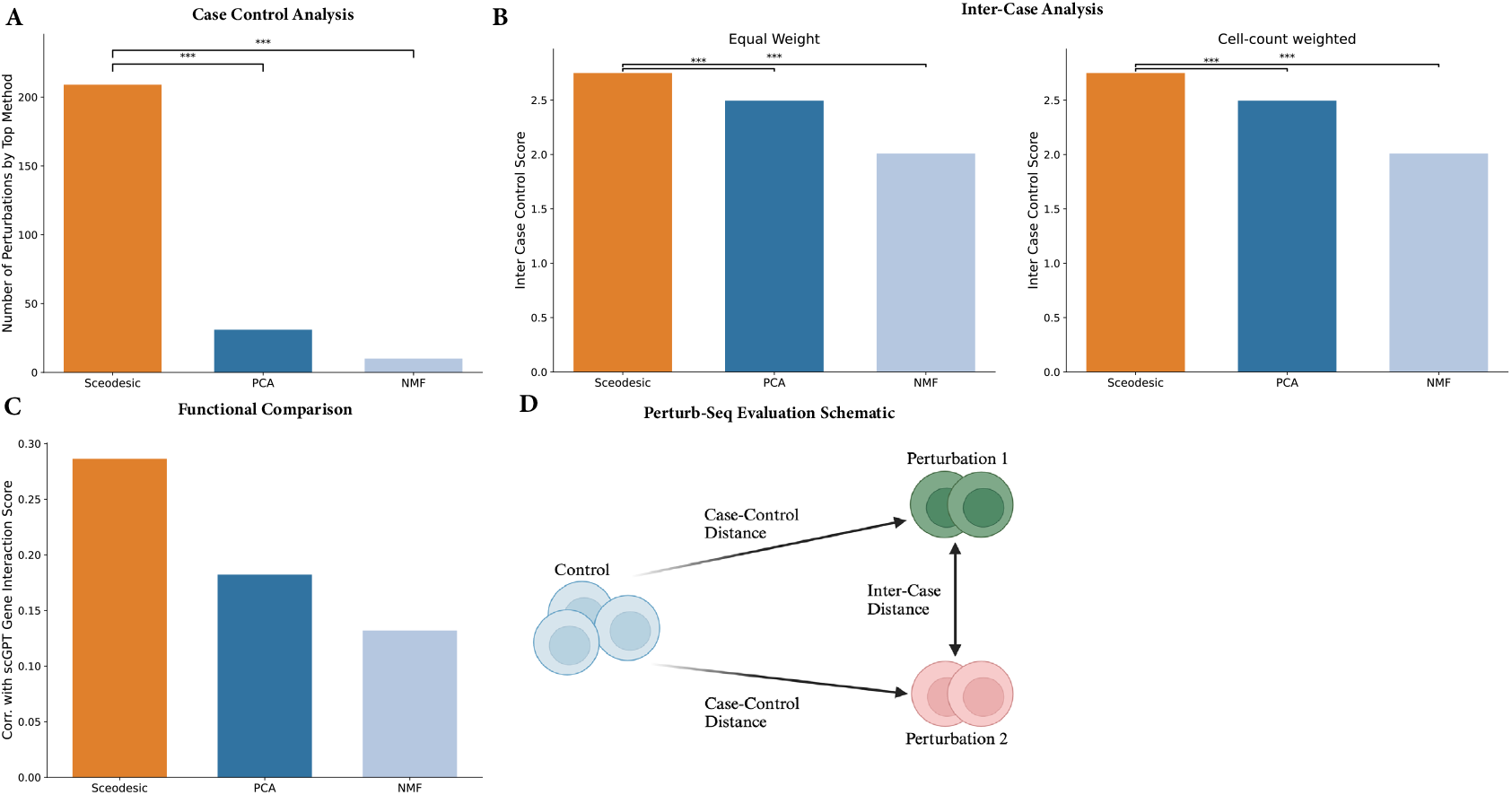
Comparing program estimation methods on Perturb-Seq data. **A)** Number of distinct perturbations for which each method has the highest case–control score. **B)** case–control score for each method, averaged over all perturbations with equal weight, as well as averaged over all perturbations proportional to number of cells belonging to a particular perturbation. **C)** Spearman correlation of perturbation to perturbation distances with gene-gene interaction scores generated by scGPT. **D)** Biologically meaningful and discriminative metrics are determined by the ability to recapitulate transcriptional state such that a control state is further from a perturbed state than sets of cells within the control state are to each other. Similarly, different perturbations should be further from each other than transcriptional states of cells within those same perturbations. Perturb-seq facilitates such analysis of the discriminative power of a program estimation method. The case–control score, or, the ratio of case–control distance to intra-control distance, captures this notion.

**Figure 5:**
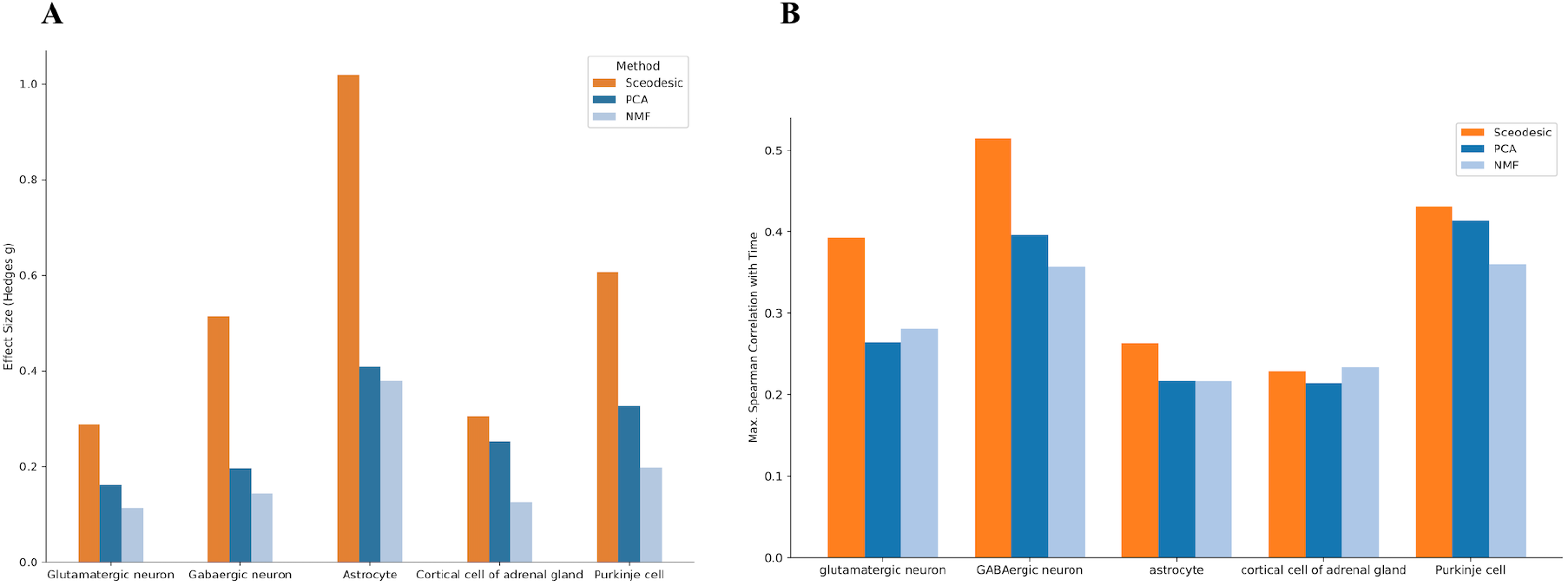
Comparing program estimation methods on time-course data. **A)** Absolute value of Hedges’ *g* effect size for change in program expression level from earliest time point to latest time point available. **B)** Spearman correlation between developmental time and program expression for the most correlated program from each method.

## Results

### The log-Euclidean metric: An expressive and mathematically consistent metric for coexpression

The practice of comparing distinct transcriptional states by taking average gene expression vectors and computing their Euclidean distance has been ubiquitous in the field of single-cell analysis, appearing everywhere from differential expression analysis to clustering and pseudotime inference. However, as datasets grow larger and more heterogeneous, distinct cell states may have similar average gene expression and thus differences in gene coexpression become necessary to discriminate between those states.

Existing work has analyzed differences in covariance across cell states using Bayesian methods or metrics that fail to respect curvature of the coexpression manifold. There is a recognition in the field that this failure may lead to spurious estimates of differential coexpression. Micheletti et al. sought to correct for these errors in the context of batch integration [2]. Furthermore, Saha et al. have used sample-specific coexpression matrices to recognize variation across individual patients [3]. Finally, Hie et al used Frobenius norms to compute distances between covariance matrices [4]. The central issue with current methods for comparing covariance across conditions is that they ignore the defining structural property that make covariance matrices what they are: their Positive Symmetric Definite property. Failing to respect the PSD property is equivalent to ignoring the curvature inherent to the space of covariance matrices. Respecting the curvature of this manifold is *necessary* for distance between objects on the manifold to be meaningful, interpretable, and consistent.

As a mathematically consistent and computationally efficient notion of distance between covariance matrices, we introduce the log-Euclidean metric. The log-Euclidean metric is a core mathematical concept from the field of differential geometry, which deals with objects that live in curved spaces. Our methodological advance rests on two key conceptual insights. The first is mathematical: Under certain conditions, the log-Euclidean metric provides the *best* possible linear approximation to the true or geodesic distance between two objects in a curved space. Covariance matrices are one such object. Therefore, we exploit the structure of the logarithmic map to approximate geodesic, distance between covariance matrices in the space of Positive Semi-Definite matrices. Furthermore, since the log-Euclidean metric is empirical, it does not impose distributional assumptions or require difficult-to-compute conjugate priors like the Wishart distribution. The second conceptual insight is computational: the matrix logarithm for covariance matrices can be computed directly from an eigendecomposition, which is exactly the same kind of analysis performed in principal component analysis. Specifically, the matrix logarithm of a covariance matrix has the same eigenvectors as the original matrix and its eigenvalues are the logarithms of the original. The matrix logarithm then follows as the Euclidean distance between the logarithms of two covariance matrices (Fig. 1D). In computing the log-Euclidean metric, we therefore deploy existing and efficient computational techniques for eigendecomposition of cell cohorts, while endowing the log-Euclidean metric with an intuitive, PCA-like interpretation.

### The log-Euclidean metric captures differences in cell state

#### Capturing differences in data simulated from artificial gene regulatory networks

We first sought to assess if the log-Euclidean metric could accurately pinpoint transcriptional similarities on simulated cell types for which ground-truth regulatory network structure is known. Using SERGIO [5], we simulated two cell types (A and B) characterized by distinct coexpression patterns that nonetheless had similar *average* gene expression across all regulated genes (Methods). This was achieved by setting high levels of expression for distinct master regulator genes corresponding to each cell type. Each master regulator was wired to induce high-coexpression between distinct pairs of genes. As a result, each cell type had similar average gene expression, while differentially wired master regulators ensured cells simulated from each cell type had different gene-gene coexpression patterns (Fig. S1).

We evaluated and compared three metrics on the simulated data: (1) *d*_*µ*_: the Euclidean distance between mean expression vectors of each cell type. (2) *d*_Σ_: the Euclidean distance between covariance matrices of each cell type. (3) *d*_log(Σ)_: the log-Euclidean distance between covariance matrices of each cell type. We scored each measure of transcriptional distance by its ability to assign a smaller distance to two cell cohorts from the same cell type (intra-cell type distance) than to two cohorts from different cell types (inter-cell type distance) [1]. The score, which we refer to as the ‘case–control score’, is the ratio of the intra-cell type distance to the inter-cell type distance. Additionally, to assess the robustness of each metric in discerning distinct cell types to the sort of technical noise frequently present in real world scRNA-seq data, we performed the same evaluations under outlier and dropout noise supported natively by the SERGIO package (Fig. S1). Outlier noise randomly increases simulated transcript counts, and is motivated by the observation that a small set of genes often have exceptionally high expression measurements in real scRNA-seq data [5]. In contrast, dropout noise randomly drops transcript counts to zero and is motivated by the fact that a high proportion of genes measured in the scRNA-seq assay frequently exhibit zero expression in any given cell [5]. To further assess robustness of the log-Euclidean metric’s performance, we simulated the noisy transcriptional data with Hill coefficients ranging from 1− 3, with the size of the Hill coefficient inducing proportional non-linearity in the regulator-gene relationships.

The log-Euclidean metric had the highest case–control scores across all conditions, remaining above 60. In contrast, the other metrics were less consistent, generating substantially lower scores in the range of 20-40 for the base case and dropout noise. Thus we confirmed that the log-Euclidean metric is not only mathematically consistent but also empirically robust to noise and regulatory non-linearities that are frequently encountered in real-world scRNA-seq data (Fig. 2D).

#### The log-Euclidean metric effectively relates transcriptional similarity to phenotypic similarity

We reasoned that if the log-Euclidean metric is effective, it should recapitulate expert cell type annotations. We therefore focused on four large scRNA-seq datasets profiling diverse organ systems and origins from the CELLxGENE portal [6]. The datasets include: the Human Lung Cell Atlas (2.3 million cells [7]), the one-million cell survey of embryonic development from Cao et al. [8], the Human Breast Cell Atlas (800,000 cells [9]), and the DCM/ACM Heart Cell Atlas (881,000 cells [10]).

For each dataset, we applied k-means clustering to compute cell cohorts and marked each cohort with its dominant cell type label. Individual manual annotations can themselves be inaccurate, and to mitigate this, we limited our analysis to cell cohorts with a dominant cell type. We reasoned that these cohorts represented regions of the transcriptional landscape where experts were most confident in their annotations. We then computed distances between all pairs of cohorts using each of the three metrics (*d*_*µ*_, *d*_Σ_, *d*_log(Σ)_). We subsequently evaluated each metric’s ability to predict label similarity. Intuitively, a good metric will assign a small distance between cell cohorts with the same labels (Methods). We used a 1-D discriminant as the classifier (i.e. true if distance *<* threshold, false otherwise), computing the area under the receiver operating characteristic (AUROC) for each metric.

Among the two second-moment (i.e., covariance-based) metrics, the log-Euclidean metric *d*_*log*(Σ)_ consistently outperformed the Euclidean metric *d*_Σ_ by 0.01 to 0.14 AUROC (4% − 8% improvement). Remarkably, the Riemannian metric also outperformed the first moment metric *d*_*µ*_, sometimes by substantial amounts (Fig. 2A). These findings demonstrated that a mathematically consistent treatment of gene coexpression contains meaningful biological information. This is notable because Euclidean differences of gene expression values (i.e. the first moment) underpin much of the extant single-cell RNA-seq analysis workflow—from PCA and NMF to mean-squared error loss in deep-learning architectures. Our finding therefore suggests that exploiting gene coexpression in a mathematically consistent way can unlock deeper insights from transcriptional readouts. However, a potential critique of our analysis is that the second moment may simply recapitulate the biological information encoded in the first moment. This stems from the fact that gene expression is often modeled as having a Negative Binomial or Poisson distribution, in which the means and variance are tightly related. However, we reasoned that this was unlikely as *d*_log(Σ)_ outperformed *d*_*µ*_ in most cases, rather than simply matching the latter’s performance. Nonetheless, we directly tested for and confirmed that the two metrics were intrinsically different.

We investigated the relative informativeness of the first moment metric and the log-Euclidean metric by exploring regimes where their agreement was high or low. For pairs of cell cohorts, we compared the two metrics and grouped these pairs by how similar they were on the first moment metric. Among cohort pairs in the bottom 25% of *d*_*µ*_, the metrics exhibit good agreement (Spearman correlation 0.54). This suggests that when the cell cohorts are highly similar transcriptionally, both metrics capture similar relationships. However, on moderately similar cohorts, we found this to no longer be the case. On cohort pairs where the first-moment differences are in the 25-75th percentile of *d*_*µ*_, the two metrics showed little agreement (Fig. 2C, Spearman correlation 0.10), suggesting that they capture different information in this regime. We posited that the greater biological fidelity of the log-Euclidean metric may stem from it preserving its performance in this regime. Indeed, we found that the log-Euclidean metric continued to fare better at capturing biological similarity between cohorts than does the distance between means, as measured by AUROC. Thus, the log-Euclidean metric captures distinct information that renders it better-suited to distinguish cell states than the Euclidean metric.

Our findings illustrate the value of considering the Riemannian manifold of gene coexpression matrices to delineate biological similarity more accurately. In turn, the performance of the log-Euclidean metric serves as rationale for computing and comparing local estimates of gene–gene covariance. However, for the sake of interpretability, these local estimates of covariance must be commensurable across cell cohorts and thus resolvable into a common set of gene programs. This motivates Sceodesic, which uses the solid mathematical foundation of the log-Euclidean metric to produce a single set of gene programs that effectively recapitulate disparate cell states.

### Sceodesic: an algorithm for identifying statistically principled and interpretable gene programs

Although the preceding experiments establish that the log-Euclidean metric is an effective and accurate measure of transcriptional distance, gaining deeper biological insight requires identifying the precise gene programs whose expression across cell states best explains their transcriptional difference. This motivates the introduction of Sceodesic, which leverages the log-Euclidean metric to resolve coexpression across disparate cell states into a common set of gene programs and associates program expression levels.

Sceodesic begins by forming cell cohorts using any unsupervised clustering method (e.g. Gaussian mixture models or k-means) that effectively groups cells into cohorts that capture local regions of the transcriptional landscape (Methods). Moreover, in cases where prior biological information such as cell membership in broad cell types or batches is known, Sceodesic can leverage such user-specified cohorts for downstream program estimation. Next, covariance matrices and subsequent eigendecompositions are computed for each cohort, generating a set of cohort-level programs or eigenvectors. Then, these local eigenvectors serve as the basis for performing a Population Value Decomposition (PVD), a technique for estimating shared eigenbases across covariance matrices. PVD compresses the union of these local programs into a smaller, single set (Fig. 1D). This is the final set of gene programs Sceodesic generates. Next, for each local covariance matrix, we estimate the optimal loadings on these study-wide programs under the log-Euclidean metric. These loadings constitute the gene program expression of cells in the corresponding cohort.

Unlike other methods for gene program estimation, Sceodesic explicitly leverages local transcriptional variability, building its common set of programs in a bottom up fashion rather than from the study-wide level. Not only are Sceodesic programs more informative regarding local cell state, but Sceodesic only requires a cell–by–gene expression matrix, rendering it as easily usable and broadly applicable as principle components analysis. By aggregating information in a local-to-global fashion, and focusing on coexpression in a mathematically consistent way, Sceodesic infers programs that accurately recapitulate the data in a principled fashion and capture significant biological variability.

#### Sceodesic programs enable parsimonious cross-modality inference

If Sceodesic is indeed effective at translating raw transcriptional readouts to phenotypic signatures, Sceodesic-based program estimates should recapitulate independent modalities or annotations on the same cells. Simultaneously-assayed multi-modal datasets provide a useful testbed for this hypothesis. Accordingly, we evaluated how well the programs generated by each method enable prediction of protein expression levels from transcriptional state using only a small number of programs.

Given previous work has shown that the best predictions of protein levels are obtained by considering sets of genes (rather than just the one gene corresponding to the protein), we benchmarked Sceodesic against two other linear approaches, PCA and NMF [11]. Linear approaches like scLinear have outperformed more complex, machine learning based approaches to predict single-cell protein abundance from RNA expression data [11]. Therefore, we limited our analysis to linear approaches that generate interpretable sets of gene programs. We considered CITE-seq data, which simultaneously measures RNA transcript abundance and marker protein abundance at the single-cell level [12]. From Hao et al.’s study of peripheral blood mononuclear cells (PBMCs) [13], we obtained CITE-seq data on 211,000 cells, comprising whole-transcriptome gene expression and protein expression levels (ADT) for 228 marker proteins. We then sought to predict protein expression levels using estimated transcriptional signatures.

We reasoned that if the programs were recognizing relevant biological functions, then a small number of programs should be able to accurately predict protein abundance. Such sparse prediction would also serve to better guide subsequent biological investigations. We estimated Sceodesic, PCA, and NMF-based programs from the transcriptional data (Methods). We next fitted lasso-regularized regression models to predict protein levels in the training data. We calibrated regularization thresholds to select the same sparsity level (10 components) for each method. Finally, we measured the performance of these sparse models on the test set, quantified as the R-squared in recovering protein expression values from transcriptional readouts (Fig. 3A).

Among the various methods, Sceodesic was the best performing model on 60% of proteins, measured in terms of test-set *R*^2^ values. In direct comparisons, Sceodesic significantly outperformed PCA (85% of cases, *p*-value =1.90 × 10^−19^ from the paired-sample t-test), and NMF (76% of cases, *p*-value =1.99 × 10^−12^ from the paired-sample t-test). In other words, a small number of Sceodesic programs were best able to explain variation in surface protein expression levels, suggesting these programs are picking up on meaningful biological functionality and can be used to parsimoniously explain that same functionality.

One of the surface proteins whose expression levels Sceodesic predicts particularly well is CD305, whose associated gene is LAIR1 (Leukocyte-Associated Immunoglobulin-like Receptor 1) (Fig. 3B). LAIR1 plays a significant role in immune system regulation, particularly in the suppression of immune responses [14]. It is found on the surface of various immune cells (T cells, B cells, and natural killer (NK) cells) where it helps modulate their activation. Recently, it has been shown that LAIR1 expression levels can be diagnostic of cancer as well as a prospective target for cancer treatment [15]. The highly predictive components that Sceodesic produces make it easier to interpret and thus experimentally investigate the functional relationships suggested by regression results (Fig. 3).

#### Sceodesic faithfully represents the transcriptional effect of perturbations

If Sceodesic truly produces meaningful quantitative representations of the transcriptional signatures of distinct phenotypes, it should effectively separate wild-type from perturbed cell-states and group cell-states resulting from similar perturbations. This motivates using Perturb-seq data to test the biological accuracy of Sceodesic by considering whether: i) cells in the wild-type group are closer to each other than to those in a perturbed state; ii) cells in distinct perturbation states are further from each other than to cells in the same perturbation state; iii) cells administered functionally-similar perturbations are closer to each other than to cells administered functionally-distinct perturbations. Repogle et al. conducted a genome-scale Perturb-seq that targeted 9866 genes using CRISPRi knockdowns on 2.5 million human cells [16]. We analyzed 2 million gene expression profiles belonging to cells in the CML K562 cell line, which are artificial antigen-presenting cells [17]. We compared Sceodesic to programs generated by PCA and NMF.

First, we evaluated the ability of Sceodesic, NMF, and PCA to distinguish perturbation states from the wild-type or control state. In Perturb-seq, control cells are characterized by non-targeting sgRNAs, which do not target any particular site on the genome. Following Ji et al. [1] we computed *intra–control* distances between cell cohorts and compared them to inter control–perturbation distances, which we refer to as *case–control* distances. We scored each method by its ratio of the case–control distance to the intra–control distance, which acts as a measure of how well a set of gene programs differentiates distinct cell states. To compute the intra–control distance, we randomly partitioned the control state cells and computed the average difference between gene programs over random pairs of cell cohorts in this group. This was repeated and averaged over 100 partitions, to account for noise. To compute the case–control distance, we took the average gene program expression of the entire control group and computed the distance to the corresponding average profile of each perturbation. Second, to evaluate Sceodesic’s ability to differentiate distinct perturbation states from one another, we selected the 250 perturbations with the greatest number of cells. Treating each of these, in turn, as the reference set, we applied the previous approach to estimate case–control separation. Finally, to evaluate Sceodesic’s ability to further group functionally similar perturbations and separate functionally dissimilar perturbations, we computed the Spearman correlation of scGPT gene–gene embedding similarity scores to the distances between distinct perturbation states in Sceodesic, PCA, and NMF. scGPT is a foundation model for single-cell biology that leverages generative pretrained transformers on large-scale single-cell sequencing data to produce gene embeddings capturing key transcriptional patterns [18]. With fine-tuning, scGPT representations were able to recapitulate functionally related sets of genes, making scGPT gene-gene interaction scores a good benchmark for whether linear representations like Sceodesic programs captured the same information [18]. We reasoned that a negative Spearman correlation was desirable, because such a correlation implies a high interaction score and thus functional similarity corresponds to low distances in the embedding space, which it expected for functionally similar genes.

Among the three methods, Sceodesic performed best at separating distinct phenotypic states and grouping similar states (Fig. 4). First, Sceodesic best separated perturbation states from wild-type states, producing the highest case–control score for 84% of the 250 most abundant perturbations. Second, Sceodesic most effectively distinguished different perturbation states from one another, producing significantly higher case–control scores than competing methods on a per-perturbation equal weighted and cell-count weighted basis (Sceodesic vs. PCA p-val =1.84 ×10^−30^ and Sceodesic vs. NMF p-val 1.39 × 10^−44^ from paired t-tests). This suggests Sceodesic outperformance was broad-based, holding for both frequent and infrequent perturbations. Third, Sceodesic performed best at grouping functionally similar perturbations while separating functionally dissimilar perturbations, with Sceodesic perturbation–perturbation distances exhibiting 29% Spearman correlations with scGPT gene–gene interaction scores while PCA and NMF achieved only 18% and 13% correlation respectively. This demonstrates that Sceodesic better represents functionally similarity than competing methods.

Moreover, Sceodesic is able to substantially group functionally relevant pairs of gene perturbations that PCA and NMF classify as being very far away in those gene program expression spaces. Sceodesic alone placed PPP2R1A and ZBTB7A as very close in its gene program expression space. Both of these genes have been implicated in anoikis resistance in prostate cancer [19]. Dysregulation of Anoikis is a critical mechanism involved in tumor metastasis [20].

These results suggest that the Sceodesic gene program expression space resolves raw gene expression into biologically accurate and relevant programs which capture important functional characteristics present in Perturb-seq data. Sceodesic’s ability to better capture perturbational phenotypes from transcriptional readouts creates potential for better in-silico planning and prioritizing of select perturbations in future experiments. In turn, Sceodesic programs could allow for more efficient, informed traversal of the immense perturbational landscape.

#### Sceodesic program expression captures time-series variation of development processes via interpretable gene programs

A fundamental goal of developmental biology is characterizing the transcriptional drivers and markers of embryonic development. Towards that, we assessed Sceodesic’s ability to resolve gene expression into biologically meaningful programs that exhibited high correlation with a key biological covariate: time. Cao et al. collected 121 human fetal samples with single-cell RNA-sequencing amounting to 4 million cells in total. The dataset had 15 organs represented from the 10th to the 18th week of the post-fertilization stage resolved into 8 experimental time stages [8].

We estimated Sceodesic, NMF, and PCA programs on this dataset, and asked two related questions towards identifying transcriptional markers. First, are there specific gene programs that exhibit high Spearman correlation with time-course? Second, for gene programs whose expression levels were highly correlated with time, was there a substantial change in the level of program expression from the earliest available time point to the latest? The first question asks if some gene programs have a clear temporal gradient, and the second asks whether the strength of that gradient is substantial. To answer these questions, we isolated the gene program from each method that had the strongest Spearman correlation with time after sub-setting for cell type. We considered five cell types that occurred most frequently in the dataset: Glutamatergic neurons, GABAergic neurons, astrocytes, cortical cell of adrenal gland, and Purkinje cells. We also computed the effect size, as measured by Hedges’ *g*, of that program’s expression for cells in the first time period as compared to the second (we computed the absolute value of Hedges’ *g* as expression levels are arbitrary up to arithmetic sign and thus so is the effect size). All estimation was done in an unsupervised fashion, meaning no method was able to access the developmental time points or cell types associated to each cell in the dataset.

For four out of five cell types, a Sceodesic program exhibited the highest Spearman correlation with time, with the largest outperformance occurring for the two most prominent cell types in the dataset, Glutamatergic and GABAergic neurons. This fact is in line with our more general observation that larger datasets allow Sceodesic to produce gene programs that are based on more accurate local covariance estimates. Therefore these programs are more likely to capture important functional relationships between genes and the biological covariates they correspond to—in this case development. Moreover, Sceodesic programs also demonstrated the largest effect size between the first and last development time points available for each cell type, and in the case of astrocytes, the effect size of the best Sceodesic program more than doubled the next best-performing method. In the case of the Glutamatergic neuron, the Sceodesic gene program with the maximum time correlation prominently featured a set of genes that, upon gene set enrichment analysis in Enrichr [21], were significantly associated with neuron projection (Benjamini-Hochberg corrected p-value = 0.045). Neuron projections are prolongations of neurons that characterize the development of the embryonic brain and are critical for achieving widespread connectivity between neurons [22]. Therefore, Sceodesic’s programs not only pick up genes that statistically vary with time, but feature biological connections to the course of development itself for specific cell types.

## Discussion

The central promise of single-cell transcriptomics is to measure cellular heterogeneity at a resolution not previously possible. For these data to power discovery and clinical development, we need mathematical signatures of transcriptional measurements that faithfully capture the distinct cell states comprising such biological heterogeneity. By leveraging the tools of differential geometry, specifically the log-Euclidean metric, Sceodesic resolves gene expression measurements into a small number of linear, sparse and interpretable gene programs whose statistical properties display high correspondence to observed phenotypes.

We sought to validate Sceodesic against independently-measured ground truth rather than against internal consistency with computational measures such as clustering overlap with potentially-erroneous cell type annotations. Towards this, we exploited multi-modal and perturbation-based single-cell datasets where transcriptional signatures can be evaluated against real biological ground truth, in the form of measurements from other modalities like surface protein expression levels or case–control differences. These experiments confirmed that the Sceodesic programs accurately uncover the signature of cell phenotype and complex cellular activities present in transcriptional state.

Our development of Sceodesic is motivated by the advent of larger single-cell datasets. Previously, with smaller datasets, the focus has primarily been on the first moments of the data. Even when grouping cells for analysis, existing work such as the metacell approach from Baran et al. has focused on the first moment (i.e., averaging gene expression within cell states) [23]. With these small datasets, accurately estimating higher moments of the data, such as covariance, was difficult enough at the global whole-dataset scale, let alone for specific cell-states. But the availability of larger datasets where more observations cover the same cell state allows for the expression patterns of distinct cell states to be better characterized. Specifically, we focused on studying the covariance of gene expression, which naturally resolves into gene programs that serve as transcriptional signatures of complex biological phenomena and cell phenotypes. Crucially, our covariance estimates are computed locally for each cell cohort and then aggregated, side-stepping the risk of batch effects and other measurements to confound the estimates. To enable efficient and accurate estimation of interpretable programs, Sceodesic reaches into the mathematical theory of Riemannian manifolds and, specifically, the cone of positive symmetric definite matrices. While the log-Euclidean metric serves as the mathematical cornerstone that enables the efficient and correct comparison of gene coexpression across cell states, Sceodesic’s efficiency and interpretability stems from our innovative combination of this deep mathematical insight with computationally robust estimation of shared eigenbases. We additionally introduce a principled technique to address ill-conditioned covariance matrices that expands the general applicability of second-moment methods in single-cell analyses.

Sceodesic’s ability to accurately characterize perturbations and parsimoniously predict cross-modality readouts opens up many possible applications. Compared to previous work, Sceodesic is more effective at bridging the transcriptomic–proteomic divide as its linear programs consistently recapitulate proteomic readouts more accurately than existing approaches. Thus, in a basic science or diagnostic setting, Sceodesic allows for a single transcriptional assay to give substantial information about protein expression levels for hundreds of proteins, saving time and resources, as well as enabling more informed decision making about which assays should be subsequently employed. In effect, a single scRNA-seq assay can, in conjunction with Sceodesic protein predictions, serve as a panel of tests for cell state. More broadly, Sceodesic programs fit seamlessly into existing single-cell analyses. For instance, they can be used to supplement PCA in a clustering analysis, and Sceodesic gene program expression levels can play a role in computing nearest neighbor graphs that are upstream of many pseudotime, clustering, and dimensionality reduction analyses. Looking ahead, Sceodesic could be extended to include information about both the first and second moments of gene expression, generating multivariate normal representations of data. While we focus on specific scRNA-seq applications, our innovations are also relevant to other problems where coexpression is of interest–for instance, in analyzing protein abundance across different cellular cohorts. We anticipate that Sceodesic will help unlock fundamental biological insights from modern single-cell data.

## Methods

### The Sceodesic algorithm

The goal of the Sceodesic algorithm is to estimate biologically meaningful gene programs and recapitulate gene expression data as levels of gene program expression. The input to Sceodesic is a cell-by-gene count matrix *X* ∈ ℝ^*N*×*G*^. Sceodesic outputs a set of *h* programs *Û* ∈ℝ^*h*×*h*^ (i.e., each program is a *h*-dimensional vector) and a matrix of cell-by-program expression, *S*_*sceo*_ ∈ℝ^*N*×*h*^. Here, *h* is a user-specified parameter designating the number of variable genes to be considered. The computation consists of four main steps:

1. <u>Estimate cell cohort membership</u>. We delineate groups of cells by either hard clustering (K-Means or Leiden clustering) or soft-clustering (Gaussian mixture models) clustering on log-and count-normalized gene expression. Cells with similar transcriptional profiles will be grouped together in the same cell cohort.
2. <u>For each cell cohort, compute a covariance matrix</u>. We newly introduce *condition-volume normalization*, which overcomes the common problem of a singular sample covariance matrix by regularizing it into a well-conditioned, well-scaled positive-definite matrix closest to it. The eigenvectors of this matrix constitute cohort-specific gene programs.
3. Using population value decomposition, we <u>share gene program definitions across cell cohorts</u> to learn a shared eigenbasis that is applicable globally. We use sparse estimation to learn a set of *h* interpretable features that best recapitulate local, cohort-specific features across the entire dataset. These features are Sceodesic’s estimated gene programs *Û*.
4. <u>We reconstruct local cell cohort-specific coexpression matrices</u> in terms of the shared program definitions while respecting the manifold of PSD matrices. Crucially, the reconstruction error minimized is the log-Euclidean distance between the condition-volume normalized covariance matrix of each cell cohort and the estimated reconstruction. This ensures that biological information is preserved in the reconstruction step.

#### Cell cohort construction

Sceodesic constructs cell cohorts by partitioning a scRNA-seq dataset into *n* cell cohorts denoted *c*_*i*_, *i*=1,…*n* under the premise that cells alike in their transcriptional state will be grouped under the same cell cohort. In principle any single-cell clustering method can be used, but we chose GMMs because we found it to deliver reasonably balanced cohorts compared to other methods and because it induces meaningful variability in the representations of different cells. Balanced cohorts are important because they increase the probability of obtaining well-estimated local coexpression matrices, which are the base estimator of Sceodesic. Well-balanced cell cohorts avoid extremely small clusters which would need to be heavily regularized to reside on the positive-definite manifold, distorting the spectrum of the raw covariance matrix.

#### Sceodesic computes condition-number normalized covariance matrices

Next, Sceodesic generates positive-definite covariance matrices for each cell cohort. First, the data matrix *X* is restricted to columns representing only *h* variable genes (determined by a gene having high dispersion across a large proportion of clusters, Sec. A), denoted *X*_*hvg*_ ∈ ℝ^*N*×*h*^. This ensures the number of samples needed per cell cohort to avoid a degenerate covariance matrix is not prohibitive. For each cell cohort *c*_*i*_, the sample gene expression covariance matrix Σ_*i*_ ∈ ℝ^*h*×*h*^ is computed. However, for some cohorts, this matrix may have zero eigenvalues or be ill-conditioned, rendering matrix logarithms unstable. We address this by regularizing Σ_*i*_, first computing its spectral decomposition 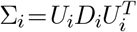, then, given a condition number *M* as the target, rounding up the smallest eigenvalues to enforce the target. Let the largest eigenvalue of *D*_*i*_ be *σ*_*max*_. We define the diagonal matrix 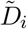 as:

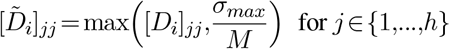

This operation is well-justified mathematically. Tanaka and Nakata showed that 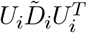 is the semi-definite matrix with condition number at least *M* that is closest (in a Frobenius-norm sense) to Σ_*i*_ [24]. Lastly, we re-scale this matrix by scaling diagonal elements of 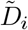 by the smallest eigenvalue 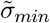, so that 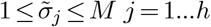. This offers two advantages: a) when mapping to the tangent space by taking logarithms, the smallest value(s) become zero, enhancing interpretability and sparsity, b) we effectively bound the determinant to be ≤*M*^*h*^. In linear algebra, the determinant has a volume interpretation, thus bounding it makes the magnitude of overall covariance comparable across cohorts, analogous to count normalization for gene expression.

#### Sceodesic learns study-wide gene programs from local gene programs via population-value decomposition

The eigenvectors *U*_*i*_ ∈ ℝ^*h*×*h*^ constitute local gene programs for each cell cohort *c*_*i*_. Sceodesic next identifies frequently-occurring and rare programs across cohorts, producing a study-wide set of programs that could be a shared dictionary globally. To do so, we adapt population value decomposition (PVD), a method first introduced in image processing [25]. PVD estimates common features between images by estimating an optimal set of vectors that serve as a common basis for reconstructing *features* specific to individual images, rather than the images *themselves*. In our context, the key insight of PVD is that the set of eigenvectors across all cohorts can itself be viewed as observations in ℝ^*h*^. Performing PCA on this set will reveal frequent and rare programs, as high- and low-ranked PCs, respectively. We select only the eigenvectors of each cell cohort sufficient to explain 90% of the variability in the cohort, in order to limit noisy features. Denoting this set 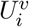, we concatenate them across cohorts,

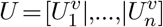

and perform a sparse principal component analysis on the rows of *U*^*T*^ as if they were themselves samples (although each row is really a gene program). This yields a series of sparse vectors *P*_*j*_ ∈ℝ^*h*^, for *j* = 1,…,*h*, that best reconstruct the features. The programs’ sparse estimation increases their interpretability, especially in context of gene set enrichment analysis. We denote *Û* as the matrix whose column vectors are *P*_*j*_, underscoring that these programs are analogous to eigenvectors.

#### Sceodesic reconstructs gene coexpression while respecting the PSD manifold

Finally, having estimated study-wide gene programs *Û*, we reconstruct cell cohort-specific covariance matrices using these programs. This amounts to finding the set of weights on the estimated programs that minimize the distance to a cohort’s covariance matrix. However, per the previous discussion, this distance computed between two matrices must respect the PSD manifold’s curvature to remain mathematically meaningful and preserve biological information. Therefore, we crucially use the log-Euclidean metric as the basis for our objective function in the reconstruction task. We note that this metric is a lower bound on the manifold geodesic distance, i.e., 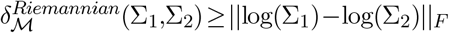[26]. The reconstruction problem for cohort *c*_*i*_ is:

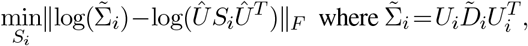

subject to the constraint that *S*_*i*_ is diagonal. Essentially, we are solving for closeness in the tangent space. Since log(*ÛS*_*i*_*Û*^*T*^)=*Û*log(*S*_*i*_)*Û*^*T*^, we work with log(*S*_*i*_), denoting it 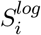. We describe the solution in Sec. A, where we derive the final form for the diagonal entries of 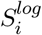:

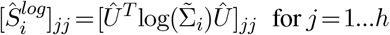

The feature weights of 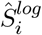 recapitulate gene program activity of cohort *c*_*i*_ on the tangent bundle of the 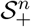 manifold, allowing for the weights to be consistently manipulated using Euclidean metrics. At the same time, though the weights correspond to shared study-wide dictionary of gene programs, the weights are themselves still specific to *c*_*i*_’s cell state.

The final product of Sceodesic is thus twofold: *Û*, a set of *h* distinct gene programs each in ℝ^*h*^, and second *S*_*sceo*_ ∈ℝ^*N*×*h*^, a matrix of cell-by-program expression. All cells in the cohort *c*_*i*_ have the feature vector 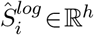. These per-cohort features can be concatenated with per-cell features from PCA or NMF to provide a representation capturing both global (PCA or NMF) and local (Sceodesic) variability.

#### Guidance on hyperparameter choices

The two key hyperparameters in Sceodesic are the number of cell cohorts *n*, and the number of variable genes *h* for which coexpression will be estimated. These parameters directly determine whether the base estimators, the sample covariance matrices of the cohorts, tend to be singular. Increasing *n* provides greater cell state granularity but decreases the average number of cells per cohort and thus increases the chances that some sample covariance matrices are singular. Similarly, a higher *h* increases the hurdle to a non-singular matrix. We recommend setting *h* to about 300, and *n* to be *N/h*, so that the typical cohort is just big enough to likely produce a non-singular covariance matrix. For a dataset of 30,000 cells, this implies *n* =100, enough to cover transient cell states in most studies. These settings are also the default in the code. There are also minor hyperparameters: the target condition number *M*, the sparsity parameter *λ* in the sparse PCA step of the population value decomposition, and the amount of explained variance *v*_*pvd*_ that determines how many cohort-specific eigenvectors should be included in the PVD. These can be left to their default values (see guidance and robustness analysis in Sec. A).

#### Validation on Simulated Data

Our main observation was that treating differences in covariance in a mathematically consistent way would allow true differences in gene program expression to be captured mathematically. A close corollary of this observation is that if two cells were known to have distinct regulatory networks governing their transcriptional state, then we would expect the log-Euclidean metric to pick up on those differences, and in particular when average gene expression was similar across cell conditions. Therefore, we created two distinct gene regulatory networks (Fig. S1).

We simulated the single cell data from these gene regulatory networks using SERGIO, and added increasing amounts of noise of several types. SERGIO takes as input a user defined gene regulatory network, in which master regulator genes are given base expression levels, and downstream genes that the GRN connects those master regulators to have their expression as some non-linear function of the master regulator genes’ expression. More specifically, SERGIO uses a stochastic differential equation (SDE) which is known as the chemical Langevin equation in simulating gene expression dynamics over time [5].

We added two forms of noise that SERGIO supports, dropouts and outliers. Dropout noise increases the probability that observed transcript counts stochastically drop to zero, despite the GRN structure staying the same. In contrast, outlier noise increases the probability that observed counts are much higher than realized in the actual network simulation.

Given this simulated gene expression data, we then computed case–control scores as described in the main text, to determine how well cell states were separated by the log-Euclidean metric, and the Euclidean metric on the mean expression vector and the covariance matrix. For all scenarios, including without technical noise, we used Hill coefficients 1, 2, and 3 which denote increasing non-linearity in the expression response of genes regulated by master regulator genes upstream in the regulatory network. For outlier noise, we used parameters: outlier probability 0.01, outlier standard deviation 0.1, outlier mean 0.5. For dropouts we used a mean parameter of 0.1 and a dropout percentile of 5% (though we ran identical experiments for percentiles of 10 and 20 percent with qualitatively similar results). For more details on SERGIO parameters and the differential equations simulated, see [5].

#### Cell type discrimination validation of distance metrics

We tested the effectiveness of the log-Euclidean metric via its ability to capture cell type annotated cell type differences. We used scRNA-seq datasets from the CELLxGENE portal [6], including Human Lung Cell Atlas (2.3 million cells [7]), the one-million cell survey of embryonic development from Cao et al. [8], the Human Breast Cell Atlas (800,000 cells [9]), and the DCM/ACM Heart Cell Atlas (881,000 cells [10]).

We ran k-means clustering to compute cell cohorts and marked each cohort with its dominant cell type label, excluding cohorts without a dominant cell type (we required cell type purity of 0.9 in terms of cell type). For the four datasets, we set *k* to keep a roughly fixed average number of cells across datasets. Therefore for the lung, fetal, breast, and heart datasets we used *k* =(500,250,200,200) cohorts, respectively in our experiments. For all datasets, we set the number of highly variable genes used in covariance calculations equal to 400. For cohort-specific covariance matrices, we used a condition control number of 50, which we have found delivers stable and accurate estimates. Computations of mean expression vectors used all *G* genes present in each dataset.

Then, we tested three metrics (*d*_*µ*_, *d*_Σ_, *d*_log(Σ)_) ability to group pairs of cohorts with the same cell type and distinguish pairs with distinct cell-types via a 1-D discriminant. We used AUROC as the performance metric. We recomputed the same AUROC performance metric for the middle 50% of observations in terms of the mean distances between clusters. To compare the intrinsic differences between the Euclidean distance of the mean vectors characterizing cohorts and the log-Euclidean metric, we used Spearman correlation.

#### Cite-seq validation experiments

To evaluate Sceodesic programs’ ability to pick up on complex biological phenomena, we used multi-modal data in which two distinct measurements were taken for the same cells. Specifically, we used data generated by Cite-Seq, which measures protein expression levels and gene expression levels at single-cell resolution. If Sceodesic programs could function as effective predictor variables for protein expression levels, then it would be good evidence the programs were picking up on transcriptional signatures of complex cellular activity. Since methods that aim to recapitulate gene expression as gene program expression can, as in the case of PCA, almost exactly capture gene expression given enough components, we wanted to test if a small number of programs were biologically informative and could compress important information in the transcriptional state. Thus, we limited each method by choosing the *k* =10 most predictive components of surface protein levels for each protein, and assessed the quality of those models.

First, for each surface protein’s expression levels, we performed variable selection by isolating each method’s most predictive programs. We did this by fitting a lasso regression with 5-fold cross validation to select the *k* =10 largest coefficients from the most predictive lasso model as the candidate programs. Next, using these selected candidate programs, we performed linear regression on those ten variables with the standard 80-20 training-test split for each protein. We assessed each model by computing the out-of-sample *R*^2^.

For Sceodesic, the hyperparameter choices were: 300 highly variable genes, 100 cohorts, 0 sparsity, 50 maximum condition number, and gamma for GMM of 0.01. For PCA, the first 300 PCs were used. For NMF, 300 components were used, with a tolerance of 1.0*e*−03 in the estimation and 300 maximum iterations.

#### Perturb-seq validation experiments

The dataset contained 1.9 million cells in total with 9867 distinct perturbations. We computed case control scores as described in Results. The case control score is a ratio between the inter-condition distance for a metric and the intra-condition distance for a metric, where the denominator is the intra-condition distance for a reference class. In this case, we used two distinct sets of reference classes. First, a proper control–we used cells with non-targeting guide RNAs as controls, which is the standard in Perturb-seq analyses. Second, we used different common perturbations as the reference class to measure the ability of metrics to separate different perturbations. For the reference class, we performed 1000 random uniform splits of the row indices into two groups and computed the Euclidean distance of the mean gene program expression in each of the two groups. For inter-condition distances, we compared mean gene program expression across those conditions using a Euclidean distance.

For PCA and NMF, we used 300 components. For Sceodesic the hyperparameters were: 300 highly variable genes, 150 cohorts, 0 sparsity, 50 maximum condition number, and *gamma* =0.001.

#### Time-course validation experiments

We computed Spearman correlations between 1. the developmental time point at which each transcriptional profile was measured for a given cell and 2. the gene program expression of a program *t* for a method *m* for each cell. We then isolated the program that exhibited the maximum Spearman time correlation for each method, and compared those. We performed GSEA using Enrichr [21], using genes whose *L*1-norm weight made up at least 95% of the *L*1-norm of the gene program vector.

For maximum effect size, we computed the absolute value of Hedges *g* between the cells of a given cell type (Glutamatergic neurons, GABAergic neurons, astrocytes,cortical cell of adrenal gland, and Purkinje cells) in the first time period they were observed in and the last time point the same cell type was observed in. The aim was to measure how significantly the gene program expression distribution changed between the start and end of the developmental process observed. Hedges g [27] is given by:

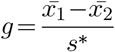

Where the denominator is the pooled standard deviation:

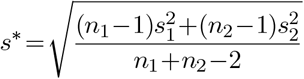

Since the program expression is arbitrary up to sign, we report the absolute value of the Hedges’ g, rather than the raw value.

#### Runtimes & memory requirement

On a dataset of 100,000 cells, Sceodesic takes approximately 20 minutes on an Intel Xeon 3GHz CPU, with runtimes increasing roughly linearly with number of cells. If interpretability of the gene programs is less crucial, these runtimes can be improved up to 3.5x by replacing sparse PCA with regular PCA during PVD. Memory requirements scaled linearly with the size of the dataset, with only 25 GB required to construct program embeddings on a dataset of approximately 1 million cells.

## Acknowledgements

We thank Alexander Wu and Kapil Devkota for helpful discussions. The authors were supported by a Duke Univ grant to RS. Some of the figures were created using Biorender. We used `GNU-Parallel` to parallelize some of our analysis.

## Declaration of Interests

None

## A Appendix

### Background

#### Additional discussion of the 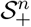 Riemannian manifold

Symmetric positive definite matrices 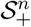 of fixed dimension *n* can be viewed as a smooth convex submanifold of the space **R**^*n*(*n*+1)*/*2^ with the inherited euclidean metric. But the direct application of this metric on 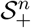 is problematic, especially because it induces finite distance between points in 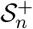 and matrices with zero or negative eigenvalues [28]. Similarly, the Frechet mean obtained using the euclidean norm induces a swelling effect, where the determinant of the euclidean norm is strictly larger than the determinants of sample points [29]. This might introduce unwanted distortions in our calculation of distance between the correlation matrices.

Many alternative norms have been developed to mitigate the finite-distance and swelling effects inherent in the Euclidean norms [28, 29]. The simplest substitute is to develop a norm in 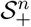 that induces isometry in the smooth map 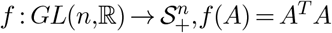[26]. The lower-bound approximation of the geodesic distance induced by this isometric norm is the logarithmic distance, given by *d*_log_(*A,B*)= ∥log(*A*) −log(*B*) ∥_*F*_, where 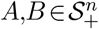 and log(*A*) represents the matrix logarithm of *A*. This *d*_log_ metric solves the finite-distance issue, as the distance between any point in 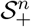 and matrices with zero or negative eigenvalues is infinity.

### Method Details

#### Selection of Locally Variable Genes

To ensure that the selection of highly variable genes is not dominated by the cells constituting larger cell cohorts, we utilize the notion of locally variable genes which allows for the selection of *h* distinct genes that are highly variable across and within cohorts, rather than only across the entire study. Consider *n* distinct cohorts *c*_1_, *c*_2_,…, *c*_*n*_ and a parameter *h* for the desired number of genes to select. Then, within each *cohort* i, we use Seurat’s HVG selection procedure (implemented in Scanpy) to obtain the top *h* genes, which we denote by the set:

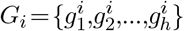

We then score each gene by the number of times it appears in a cohort-specific HVG list, i.e. for a gene *g*:

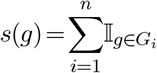

We determine the final list of locally variable genes by selecting the *h* highest scoring genes.

#### Simulated Experiment Regulatory Network Structure

Below we provide the structure of the regulatory network structure used for the SERGIO simulations. Each cell sub-type was characterized by over-expression of a single master regulator (MR) and its downstream genes. Each cell type was characterized by two distinct cell-subtypes. Edge weights denote strength and direction of regulatory relationships.

**Figure S1:**
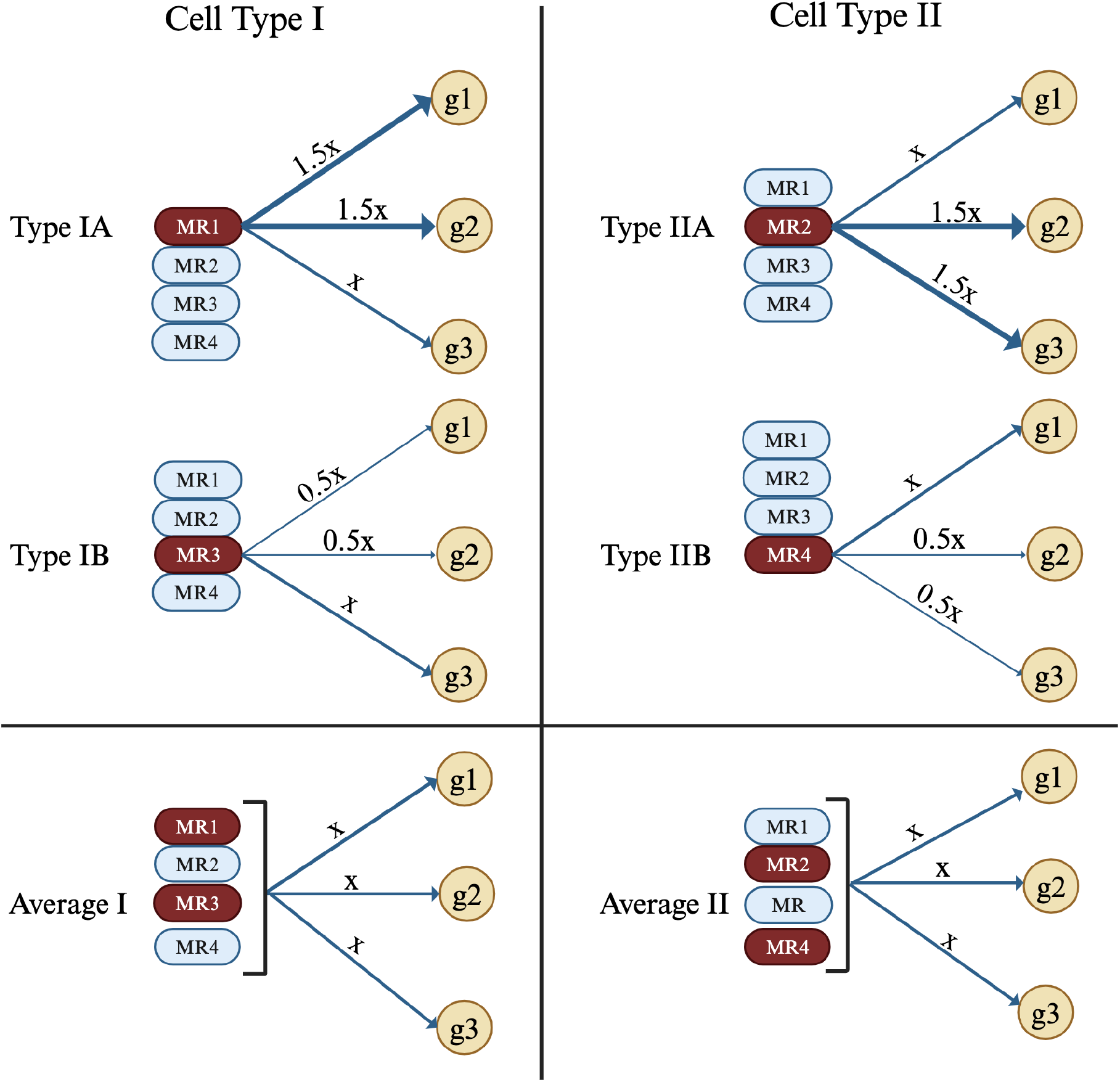
Gene regulatory network structure for each cell type in experiments on transcriptional data simulated via SERGIO, as seen in the top two entries of the grid. Each of the two cell types (I, II) is characterized by two cell sub-types. Each of the four cell sub-types (IA, IB, IIA, IIB) is generated by over-expression of a single master regulator gene (MR1-4) that picks out exactly two of the three downstream genes (g1, g2, g3) to be simultaneously over or under-regulated. Averaging over the cohorts corresponding to each cell sub-type, each cell type has identical expression of the downstream genes, as seen in the bottom two entries of the grid.

##### Proof of the optimality of 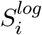

We now show that the solution *Ŝ*_*i*_ corresponding to

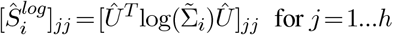

is the optimal feasible solution to the covariance matrix reconstruction problem:

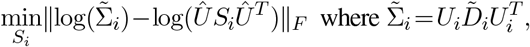

subject to the constraint that *S*_*i*_ is diagonal. The objective function of interest, which minimizes the distance from the reconstruction matrix *ÛS*_*i*_*Û*^*T*^ to the condition-normalized sample covariance matrix 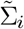 under the log-Euclidean metric, is equivalent to:

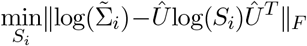

Factoring on the left and right yields:

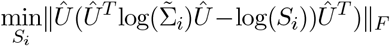

But since the columns of *Û* are orthonormal and do not change the norm of any matrix, the preceding minimization is equivalent to minimizing the simpler objective:

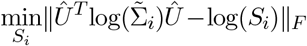

Setting 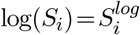, we have:

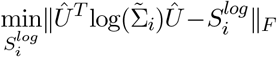

But since 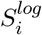 must be diagonal because *S*_*i*_ must be diagonal, the optimal feasible solution is just the diagonal of the first term 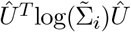, in other words:

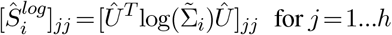

This concludes the proof. In the context of our method, we note that this solution is exact when *λ* =0 i.e. when regular PCA is used in the population value decomposition.

##### The difference of two Hermitian matrices with same eigenvectors is determined by their eigenvalues

For two Hermitian matrix ***A*** and ***B*** with common eigenvectors ***U***, the Frobeinus norm of their difference (or ∥***A***−***B***∥_*F*_) equals the *ℓ*^2^ norm of difference between their eigenvalues.

*Proof:* ***A*** and ***B*** can be both be diagonalized by a shared basis ***U***, as ***A***=***UD***_***A***_***U***\* and ***B*** =***UD***_***B***_***U***\* (where ***U*** is unitary and ***D***_***A***_,***D***_***B***_ are diagonal matrices containing eigenvalues of ***A*** and ***B*** respectively). Now, the equivalent definition of ∥·∥_*F*_ is

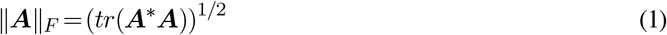

Simplifying ∥***A***−***B***∥_*F*_ using (1), we get

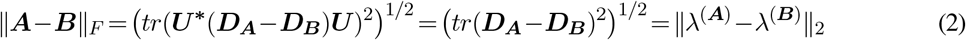

Where, *λ*^(***A***)^ and *λ*^(***B***)^ are the eigenvalues of ***A*** and ***B*** corresponding to the unitary matrix ***U***

#### Cell cohort construction

Sceodesic constructs cell cohorts by partitioning a scRNA-seq dataset into n cell cohorts ci, i=1,…n under the premise that cells alike in terms of their transcriptional state will be grouped under the same cell cohort. In principle any single-cell clustering method can be used, but we chose Gaussian mixture models because we found this method to deliver reasonably balanced clusters, which is important because well-estimated coexpression matrices of cell cohorts are the ‘base estimator’ of Sceodesic. Well-balanced cell cohorts avoid extremely small clusters which would need to be heavily regularized to reside on the positive-definite manifold, distorting the spectrum of the raw covariance matrix. We describe GMMs and soft-clustering below.

To account for the noisy nature of scRNA-seq data leading to uncertainty in clustering, we took a soft-clustering approach inspired by the classic Gaussian Mixture Models (GMM). The GMM model postulates the following data-generating mechanism: cell *i* is drawn from a mixture of *K* components (latent variable *z*_*i*_; scRNA-seq measurements of *G* genes 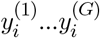 for cell *i* is given

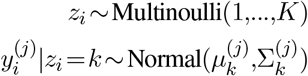

Here, 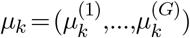 is the mean expression vector for component *k*, and Σ_*k*_ is a *G*×*G*. In our model, we constrain Σ_*k*_ to be a diagonal matrix where all mixture components have equal variance; that is, Σ_*k*_ =*σ*^2^*I* for some scalar *σ*^2^. In this case, the GMM model reduces to the classic *K-Means Clustering* model, with soft assignments.

To compute responsibilities for the soft clustering, we use the classing *K-Means* algorithm for computing cluster centers 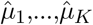, where 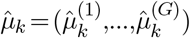. The responsibility for cell *i* belonging in cluster *k*, denoted *r*_*ik*_, is computed according to the Gaussian Mixture probability model:

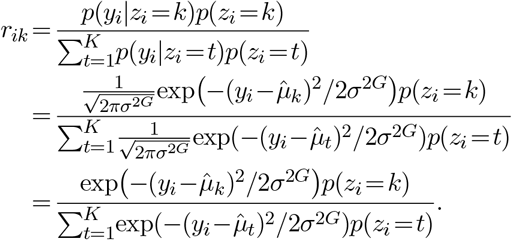

To simplify computations, we treat *σ*^2^ as a hyperparameter, denoting it as *γ*. We tested a range of values for *γ*, and found *γ* =0.01 to perform reasonably well (should we show experiments for this?).

To speed up computation, we observed that cluster responsibilities were minuscule for clusters other than those corresponding to the 5 closest centroids. To speed up computation, therefore, for each cell *i* we defined a set of clusters *C*_*i*_, consisting of the 5 clusters whose centers are closest to the observed gene expression *y*_*i*_. (For example, if the 5 “nearest” clusters to cell *i* are 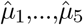, then 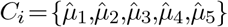). We then assumed responsibilities for clusters outside of *C*_*i*_ to be zero, and a uniform marginal distribution for the latent variable *z*_*i*_ within each cluster. The cluster responsibility for this simplified assignment scheme then becomes

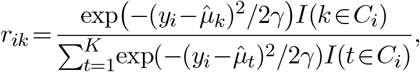

where *I* is the indicator function. We define the responsibility matrix

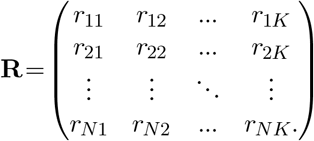

Given the embedding matrix

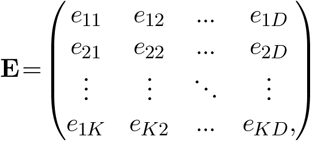

we then compute the soft embeddings assignment for the cells as a single matrix multiplication:

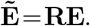

## Notes

https://singhlab.net/sceodesic

### Competing Interest Statement

The authors have declared no competing interest.

### Summary of Updates

Additional analysis with more datasets. The whole manuscript was edited as well.

